# Localized nanoscale induction by single domain magnetic particles

**DOI:** 10.1101/2020.07.16.207126

**Authors:** Michael G. Christiansen, Nima Mirkhani, William Hornslien, Simone Schuerle

## Abstract

Single domain magnetic nanoparticles are increasingly investigated as actuators of biological and chemical processes that respond to externally applied magnetic fields. Although their localized effects are frequently attributed to nanoscale heating, recent experimental evidence casts doubt on the existence of nanoscale temperature gradients in these systems. Here, using the stochastic Landau-Lifshitz-Gilbert equation and finite element modelling, we critically examine an alternative hypothesis that localized effects may be mediated by the induced electric fields arising from the detailed dynamical behavior of individual single domain magnetic particles. We apply our model to two significant case studies of magnetic nanoparticles in alternating magnetic fields: 1) magnetogenetic stimulation of channel proteins associated with ferritin and 2) catalytic enhancement of electrochemical hydrolysis. For the first case, while the local electric fields that ferritin generates are shown to be insufficient to perturb the transmembrane potential, fields on the surface of its mineral core on the order of 10^2^ to 10^3^ V/m may play a role in mass transport or release of iron ions that indirectly lead to stimulation. For the second case, our model indicates electric fields of approximately 300 V/m on the surface of the catalytic particles, with the highest interfacial electric field strengths expected during reversal events. This suggests that the nanoparticles best suited for hysteresis heating would also act as intermittent sources of localized induced electric fields in response to an alternating applied field. Finally, we put the magnitude and timescale of these electric fields in the context of technologically relevant phenomena, showing that they are generally weaker and faster.

**Popular Summary:** The possibility of using magnetic fields to exert wireless control over biological or chemical processes has stimulated vigorous research efforts across disciplines. Magnetic nanoparticles exposed to alternating magnetic fields have repeatedly been found to exert an influence at the nanoscale, for instance triggering biological responses or regulating chemical catalysis. While these effects have been attributed to nanoscale heating, recent experiments have shown that the temperature in the vicinity of magnetic nanoparticles may not differ appreciably from their surroundings. Could another nanoscale phenomenon be at work?

Here, we critically examined the idea that electric fields induced in the immediate vicinity of magnetic nanoparticles might help explain nanoscale effects. The fact that magnetic nanoparticles thermally fluctuate is widely appreciated, but the process that dominates the generation of electric fields is the rapid (typically > 1 GHz) precession that the magnetic moment undergoes during reversal events. Combining a model of the detailed motion of a single magnetic moment with numerical calculation of the induced electric field, we consider the possible role of induced electric fields in two technologically important cases. The first is stimulation of neurons with weakly magnetic ferritin and the second is enhancement of hydrogen production by catalytic magnetic nanoparticles.

Understanding the mechanism by which magnetic nanoparticles act on their surroundings is crucial to designing more optimal materials for triggering chemical and biological processes. The role of electric fields explored here also suggests the possibility of pairing magnetic nanoparticles with resonant stimuli to directly drive precession.

## I. INTRODUCTION

Single domain magnetic nanoparticles (MNPs) are increasingly investigated as nanoscale actuators that enable wireless control or enhancement of biological and chemical processes in a variety of applied contexts. Commonly, these effects have been attributed to hysteretic heat dissipation, with quasimagnetostatic alternating magnetic fields (AMF) in the kilohertz or megahertz frequency range serving to couple these nanoparticles to an external circuit. The abundance of biomedical applications employing this concept reflects the underlying appeal of wireless actuation as a basis for minimally invasive manipulation. For example, MNPs exposed to AMFs have been demonstrated to modulate the anticancer activity of nanozymes [1, 2], trigger release of liposomal cargo [3–5], and to influence the composition of the protein corona formed in vivo [6]. More controversially, a broad set of techniques known as magnetogenetics has sought to genetically impart magnetic control over cellular activity, usually by expressing the Fe storage protein ferritin in association with heat sensitive ion channels [7, 8]. Apart from medicine, the use of nanoscale effects of MNPs in AMFs has been suggested for purposes such as regulating catalytic activity of enzymes [9] or enhancing the electrochemical production of hydrogen [10].

Despite the widening application space of MNPs, serious and intriguing questions can be raised regarding the exact nature of the phenomena that underpin their nanoscale actuation effects. Whereas the heating of a millimeter scale droplet of a concentrated suspension of MNPs is an unsurprising consequence of thermodynamics and bulk heat transport, a nanoscale temperature increase in aqueous solutions surrounding isolated MNPs is not predicted by applicable heat transport models [11]. Recent experiments employing rigorous controls in search of evidence for nanoscale temperature gradients generated by magnetic nanoparticles have yielded striking null results [12, 13]. On the other hand, similar experiments with plasmonic gold nanoparticles of comparable size that dissipate substantially more heat per particle have led to observations of surface temperature that are in reasonable agreement with bulk heat transport [14, 15]. While the possibility of effects such as intermittent transient heat spikes cannot be ruled out entirely [13], it is becoming increasingly clear that physical processes other than heating may be required to explain nanoscale actuation mediated by MNPs.

Here, we critically examine the possibility that localized induced electric fields may play some role in explaining effects previously attributed to nanoscale heating. This possibility does not appear to have been seriously considered, perhaps due to the small scale of MNPs or because modelling hysteresis typically averages reversal behavior over large ensembles of MNPs, allowing the detailed dynamical behavior of individual magnetic moments to be neglected. Nevertheless, the magnetization of MNPs has long been understood to rapidly precess during coherent reversal and thermal fluctuation [16], a phenomenon that can be expected to dominate the locally induced electric fields. Using the stochastic Landau-Lifshitz-Gilbert equation (sLLG) to predict the dynamical behavior of MNPs, we examine localized induced electric fields, focusing on two important cases (Fig. 1). In the first, we examine possible roles for induced electric fields in ferritin-based magnetogenetics. Despite its weak magnetic moment of about 300 *μ_B_*, ferritin is predicted to produce surprisingly high electric field magnitudes in its immediate vicinity. Appreciable interaction with the gating charges of ion channel proteins can be ruled out, yet electric fields at the surface of ferritin’s mineral core may still offer a mechanistic clue linking Fe ion release and applied magnetic fields in magnetogenetics. In a second case, we consider the magnitude of electric fields occurring the surface of metallic nanoparticles that have been reported to enhance electrochemical production of hydrogen [10].

**FIG. 1.**
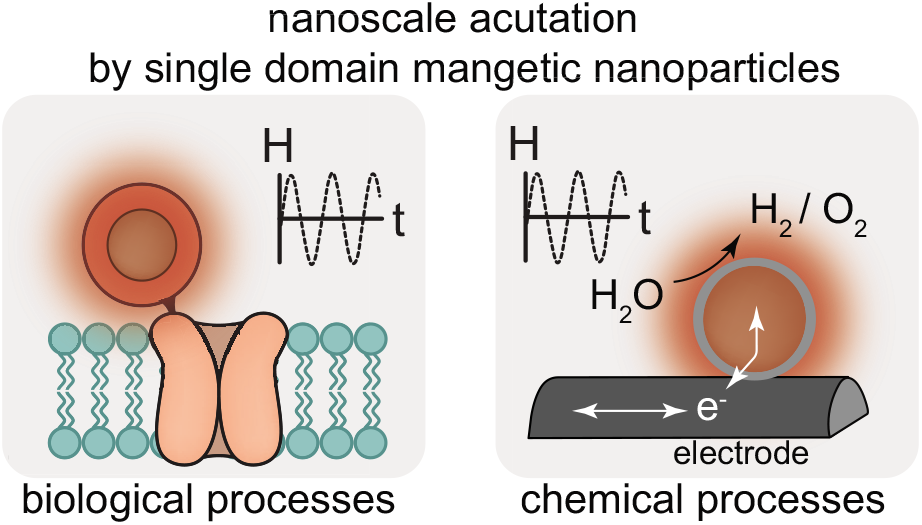
Single domain magnetic nanoparticles have increasingly been used as nanoscale actuators, coupling externally applied magnetic fields to biological processes such as the opening of channel proteins (left) or chemical processes such as catalysis (right). Although these effects are frequently attributed to nanoscale heating, the actual mechanism remains unclear.

In both case studies, the induced electric fields are surprisingly large, yet far smaller than the electric fields inherently present in the neuronal membrane or in accumulations of interfacial charge at an electrode. The rapidity of magnetic precession accounts for these unexpectedly large electric field values, yet simultaneously explains why these fields vary too quickly for capacitive effects to further magnify interfacial electric fields. We conclude by discussing possibilities for experimental investigation and the considering implications of our findings on the design of MNPs as nanoscale actuators intended to exploit these effects.

## II. MODELLING ELECTRIC FIELDS GENERATED FROM DYNAMICAL BEHAVIOR OF SUPERPARAMAGNETIC MOMENTS

To simulate the dynamical behavior of single domain magnetic particles with properties corresponding to the cases of interest, we numerically solved the stochastic Landau-Lifshitz-Gilbert equation, which describes precession of the magnetic moment of single domain particles under thermal agitation [16, 18, 19]:

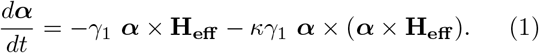

***α*** is a unit vector corresponding to the magnetization direction of the particle, *t* is time, and *κ* is a unitless damping constant. *γ*_1_ is related to *κ*, the permeability of free space *μ*_0_, and the gyromagnetic ratio *γ* as follows:

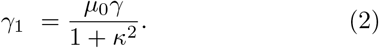

The effective field **H_eff_** is defined in terms of the overall energy function *U* of the moment *m* of a particle,

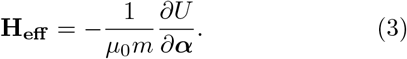

**H_eff_** can be broken into a sum of several contributions,

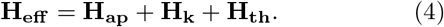

**H_ap_** is the applied field, **H_k_** is a term that captures the influence of anisotropy, and **H_th_** is the thermal field, a stochastic term. For uniaxial anisotropy, **H_k_** can be expressed in terms of the barrier energy *U_B_* separating the easy axes,

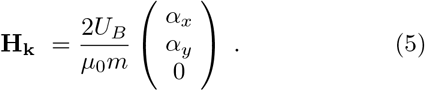

**H_th_** is defined according to statistical properties,

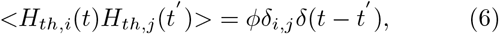

where *ϕ* is a constant proportional to *k_B_T* and inversely proportional to *m* [16]. To solve Eq. 1, we employed a fourth order Runge-Kutta algorithm to advance the system in time, renormalizing the unit vector ***α*** after each timestep. For the *i^th^* component of **H_th_**, stochastic noise was introduced by a multiplying a constant *H*_*th*0_ with a normally distributed random variate. Since the magnitude of *H*_*th*0_ depends on both physical parameters and features of the model such as the timestep, its value was determined empirically by removing anisotropy and iteratively fitting *H*_*th*0_ to produce the magnetization expected at a single point of the expected Langevin function. The result of propagating the system forward in time is precession of the magnetic moment of the single domain MNPs, subject to thermal fluctuation (Fig. 2a). Time evolution of the z projection of ***α*** (Fig. 2b) shows that the magnetic moment tends to be oriented along one of its preferred axes with occasional stochastic jumps between preferred directions, a behavior that has also been observed experimentally at lower temperatures and longer timescales [20].

**FIG. 2.**
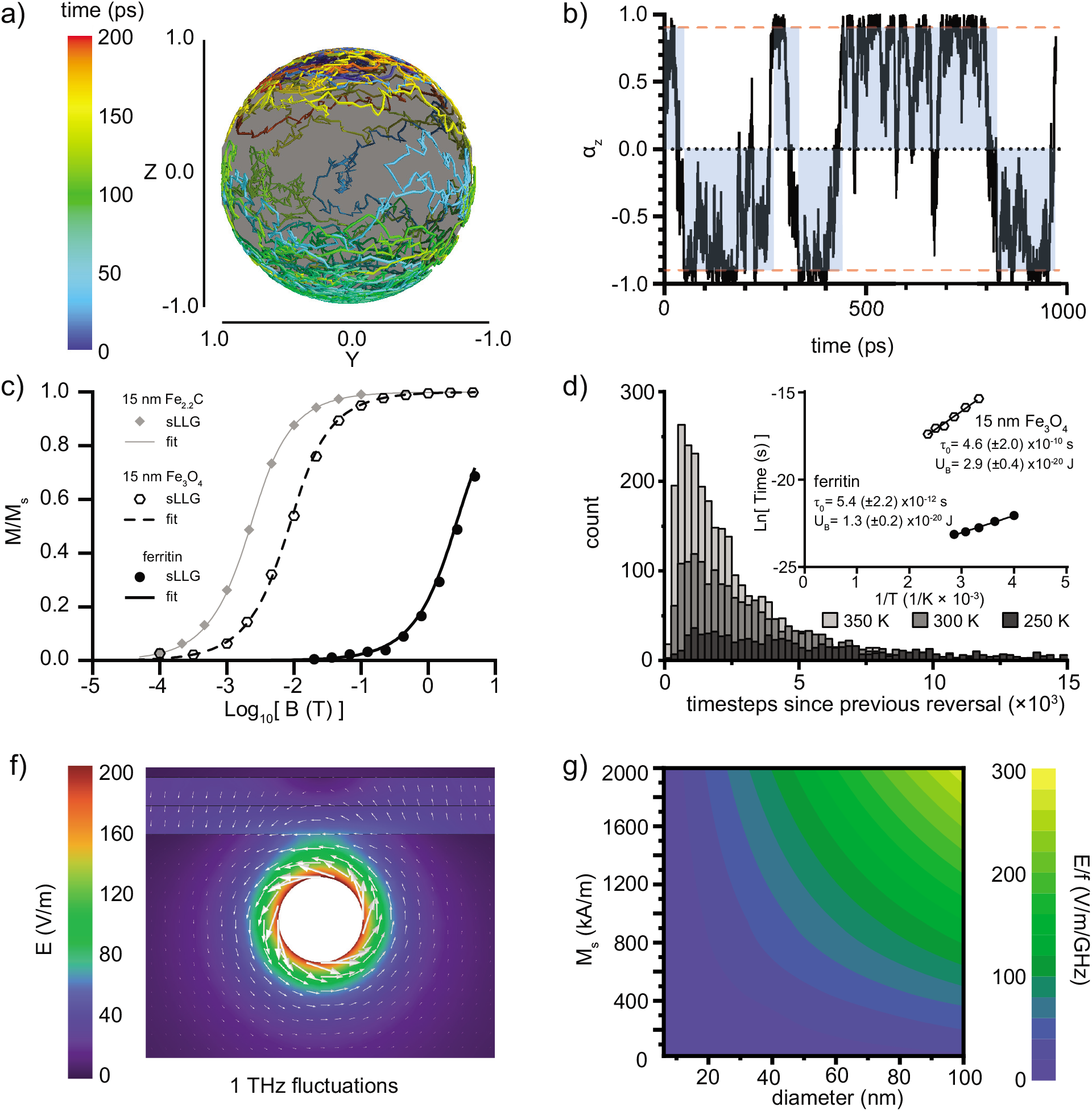
Preliminary validation of model for dynamical behavior of magnetic nanoparticles and resulting induced electric fields. a) 200 ps of the time evolution of the orientation of the magnetic moment of ferritin. Plot created with Mayavi in Python [17]. b) Z projection of the unitless magnetization of the magnetic moment of ferritin showing stochastic fluctuation over its energy barrier. c) Langevin functions shown for particles with anisotropy removed possessing the same moments as ferritin (solid line), 15nm magnetite particles (dashed line), and 15nm Fe_2.2_C (grey line). Time-averaged magnetization predicted from sLLG model versus applied field for ferritin (solid points), 15nm magnetite (open hexagons), and 15nm Fe_2.2_C (grey diamonds). Each point is the average of a simulation with 6×10^6^ to 6×10^7^ steps. d) Histogram of timesteps elapsed since previous reversal over barrier for ferritin model at three temperatures investigated by rescaling the thermal field term (250 K, 300 K, and 350 K). Mean timescale of reversal over anisotropy barriers in the absence of an applied field was extracted, with results fitted in terms of the Néel-Arrhenius model for temperature dependence of relaxation rate on temperature (inset). Fitted values are shown with uncertainty reflecting 95% confidence intervals in the linear fits. f) Induced electric fields are plotted for a ferritin moment near a membrane fluctuating at 1 THz. g) A generalized parameter space considers the magnitude of the electric field induced at the surface of hypothetical single domain particles spanning a size range from 6 to 100 nm diameter and a magnetization from 20 to 2000 kA/m. The electric field varies linearly with frequency and the values shown are normalized to 1 GHz fluctuations.

To validate the model, we assessed its convergence to the Langevin function, an analytical expression for the equilibrium behavior expected for ensembles of freely orienting magnetic moments in an applied field. By removing the contribution of anisotropy to the effective field, maintaining a consistent value for *H*_*th*0_, and calculating time-averaged magnetization versus applied field, good convergence to the Langevin function is found for the MNPs modeled in this study (Fig. 2c).

Basic descriptions of dynamical behavior of anisotropic magnetic nanoparticles typically emphasize the Néel-Arrhenius relationship, which describes temperature dependence of stochastic reversal between preferred axes over an anisotropy energy barrier [21]. In the absence of an applied field, the timescale of stochastic reversal *τ* can be expressed as a function of an attempt rate *τ*_0_, the magnitude of the intrinsic anisotropy barrier *U_B_*, the Boltzmann constant *k_B_*, and the absolute temperature *T*:

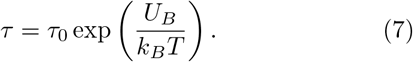

Varying the temperature by rescaling the stochastic term in the sLLG equation and extracting distributions of lifetimes between reversal events, ln *τ* versus 1/*T* can be plotted (Fig. 2d). The linearity of these curves shows convergence to expected functional behavior, and fitting these curves reproduces the values for *U_B_* are assumed by the model. The order of magnitude of the attempt rates suggested from these fits agrees well with values extracted from characterization of reconstituted ferritin [22] and the values typically assumed in modeling magnetite particles [23]. Since these were not direct input parameters for the sLLG model, it suggests that the model is capturing physical features salient to predicting dynamic behavior.

Next, to model the induced electric field arising from the changing magnetic moment of nanoparticles, Maxwell’s equations were numerically solved using the finite element method for single domain magnetic moments evolving in time.

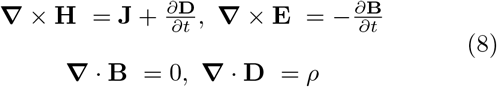

Here, **J** represents the current density, **D** is electric flux density, **E** is the electric field, and *ρ* is electric charge density. Along with the following constitutive relations, these form the set of partial differential equations that describe distribution of electric and magnetic fields in the vicinity of the nanoparticles.

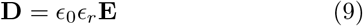

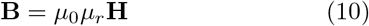

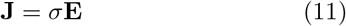

*ϵ*_0_ is permittivity of vacuum, *ϵ_r_* is the relative permittivity, *μ*_0_ is the vacuum permeability, and *μ_r_* is the relative permeability. *σ* is the electrical conductivity.

For the magnetogenetics case study described in Section III, a simplified 3D geometry of a nanoparticle located at the cytoplasmic side of the plasma membrane was reconstructed in COMSOL Multiphysics, where the Magnetic Fields interface was employed as the numerical solver. The computational domain consisted of four subdomains representing cytoplasm, nanoparticle, plasma membrane, and extracellular medium. Frequency-dependent conductivities and relative permittivities were assigned to each subdomain, as explained in greater detail in Supplementary Section S1.

Under the predicted dynamical behavior of the magnetic moment of the nanoparticle, the set of coupled PDEs was solved for magnetic vector potential, from which the induced electric field was extracted using Faraday’s law. In instances where harmonic oscillation of the magnetic moment at fixed frequencies was considered, analysis was conducted in the frequency domain. A 100-ps time-dependent study was carried out by using fluctuation of the magnetic moment predicted by the sLLG model as the input to assess the transmembrane voltage as a function of time. Because the induced electric field is non-conservative, different paths were defined normal to the membrane and adjacent to the nanoparticle to measure the induced voltage *V_ind_* between two points *a* and *b* using the following path integral:

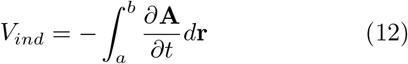

where **A** represents magnetic vector potential.

Before applying Maxwell’s equations to results of our sLLG model, an upper bound for the inductive effects of ferritin in the vicinity of a neuronal membrane were considered under the assumption of harmonic oscillation at a characteristic frequency of 1 THz (Fig. 2f). As a more general sweep of the relevant parameter space, induced electric fields were calculated at the surface of MNPs as a function of size (6 nm to 10 nm), magnetization (20 kA/m to 2000 kA/m), and frequency (1 GHz to 1 THz). Over the investigated frequency range, the peak electric field was found to scale linearly with the frequency, allowing the representation shown in Fig. 2g, in which induced electric field strength is normalized to frequency.

## III. A ROLE FOR NANOSCALE INDUCTION BY FERRITIN IN MAGNETOGENETICS?

### A. Controversy over the Mechanism of Magnetogenetic Stimulation

To exert targeted control over cellular activity in model organisms, biologists and neuroscientists have developed powerful techniques to introduce genes coding for channel proteins that, when expressed, sensitize neurons or other cell types to specific stimuli [24, 25]. Initially the preferred stimuli were visible light (“optogenetics”) or chemical substances (“chemogenetics”), however considerable interest later emerged in developing an analogous approach with magnetic fields for noninvasive actuation (“magnetogenetics”). Reports of conferred sensitivity to magnetic fields through fusion or targeting of one or more units of the Fe storage protein ferritin to a transient receptor potential vanniloid channel protein (TRPV1 or TRPV4) first appeared nearly a decade ago [7, 8], yet independent attempts to replicate these findings have often struggled to do so [26] and completely satisfactory mechanistic explanations have not been forthcoming. The presumptions that originally guided these efforts, namely that mechanical or thermal cues might act directly on TRPV channels to trigger Ca^2+^ influx, have been shown to be physically implausible [27] or to require a version of ferritin with highly unrealistic magnetic properties [28]. Due to its small magnetic moment, the interaction of ferritin with attainable magnetic fields is far too weak for mechanical actuation. Hysteresis heating of ferritin is demonstrably negligible [12], and even if extraordinary heat flow were to occur, its scale ensures that a local temperature increase is not expected [27].

Although simple physical arguments compellingly rule out direct magnetic interaction of ferritin with an external field as a mechanism for actuation, this reasoning leaves open the possibility of an indirect role for ferritin. One form this might take is for ferritin to act as a source of stochastic inductive perturbation coupled to voltage-sensitive channel proteins that sensitizes these channels to subthreshold magnetic stimuli. Because the use of rapidly varying fields to stimulate unmodified neurons in transcranial magnetic stimulation (TMS) is extensively documented [29], the role of ferritin expression in this scenario would not be to deliver the energy to reconfigure the channel proteins, but rather to generate a perturbation that effectively lowers the kinetic barrier to activation. TMS typically employs pulses with peak amplitudes greater than 1 T and *dB/dt* values on the order of 10^4^ T/s [30], although stimulation thresholds are complex, depending also on pulse characteristics [31], among other factors. Some of the reported instances of magnetogenetics involve quasimagnetostatic fields with comparatively lower amplitudes than TMS (tens of mT) that are pulsed or alternate at 100s of kHz, and their authors have suggested that time variation of the applied field may be a necessary aspect of the stimulus [32].

Another indirect mechanism of actuation by ferritin has been recently suggested to involve the magnetically triggered release of Fe^2+^ and subsequent generation of reactive oxygen species [33, 34]. Experimental evidence does appear to support this idea, showing that ferritin releases Fe content in the presence of MHz frequency fields [35] and that reactive oxygen species are implicated in magnetogenetic stimulation [33]. While these studies may help clarify the role of ferritin in magneto-genetics, the mechanism is still not yet fully satisfying from a physical perspective because it does not explain how applied fields lead to Fe^2+^ release. Here too, it is worthwhile to consider whether nanoscale induction may contribute, especially since induced electric fields are expected to be strongest at the surface of ferritin’s mineral core. In this section, we analyze the localized inductive effects predicted by a realistic model of ferritin and consider its implications on the plausibility of these two potential indirect mechanisms of stimulation for excitatory cells. Despite the remarkably high rate of change of the magnetic flux density *dB/dt* predicted in its immediate vicinity, and its ability to induce electric fields comparable to a representative magnetite particle, ultimately we conclude that the resulting electric fields generated by ferritin are insufficient to perturb nearby voltage gated ion channels. The influence of induced electric fields at the surface of the mineral core of ferritin appears to be a more relevant physical factor for explaining magneto-genetic stimulation, with a possible connection to the stimulated release of Fe^2+^.

### B. Electric field perturbation from ferritin is insufficient to directly influence channel proteins

A realistic dynamical model of the magnetic behavior of ferritin is an essential starting point for any analysis of its local inductive effects. Historically, both the reported magnetic properties of ferritin and their interpretations have varied considerably, depending on biological source and sample preparation. Nevertheless, magnetic characterization of ferritin spans nearly eight decades and can at least inform reasonable bounds for expected behavior [36]. Ferritin consists of a protein shell with an outer diameter of approximately 12 nm that surrounds a biomineralized core with diameter 5.5 to 6.0 nm in humans, ranging up to about 8nm in mollusks [37]. The size and crystallinity of human ferritin is comparable to ferritin derived from horse spleens, which is often studied. The core consists primarily of ferrihydrite with an approximate stoichiometry 5Fe_2_O_3_·9H_2_O, possibly incorporating trace phosphate impurities [38]. Whether attributable to uncompensated antiferromagnetically ordered spins or the existence of multiple phases [39], several empirical sources contend that ferritin’s ferrihydrite core is superparamagnetic at physiological temperatures with a weak magnetic moment of approximately 300 *μ_B_* [22, 40, 41]. Empirically observed effects associated with transgenic ferritin, including measurable influence on T2 relaxation in transfected yeast [42], magnetic separation in transfected bacteria [43], and electron micrographs of apparently well-crystalized cores of ferritin expressed by human embryonic kidney cells [34] support the assumption of superparamagnetic properties for ferritin in vitro or in vivo.

Approximating the moment of ferritin as a 300 *μ_B_* point dipole, the expected field magnitude at its surface is only a few mT and drops to a magnitude comparable to the geomagnetic field within 25 nm. Assuming a voltagegated ion channel with approximately 10 nm diameter (Fig. 3a upper), a ferritin directly adjacent to it should produce at most a small field of about 0.4 mT in the center of the channel. While this field magnitude is minute, inductive effects are determined by time-varying flux, and the timescale of fluctuation and precession of the moment plays a role of equal importance in determining this quantity. The effective anisotropy field *H_k_* of ferritin has been estimated from its reported blocking temperature (See Supplementary Section S2). Applying the sLLG model described in Section II to ferritin, the resulting time varying field it produces depends on position and fluctuates with frequency contributions spanning several orders of magnitude (Fig. 3b). The precession of the magnetic moment contributes high frequency components to the *dB/dt* signal, whereas stochastic reversal over the barrier tends to contribute a lower frequency component. Frequency space representations (Fig. 3c) indicate that *dB/dt* is dominated by precession, in this case peaking in the 100s of GHz. This is a high precession frequency, but a similar order of magnitude has been observed experimentally for iron oxide nanoparticles with moderate anisotropy and low magnetization values [44].

**FIG. 3.**
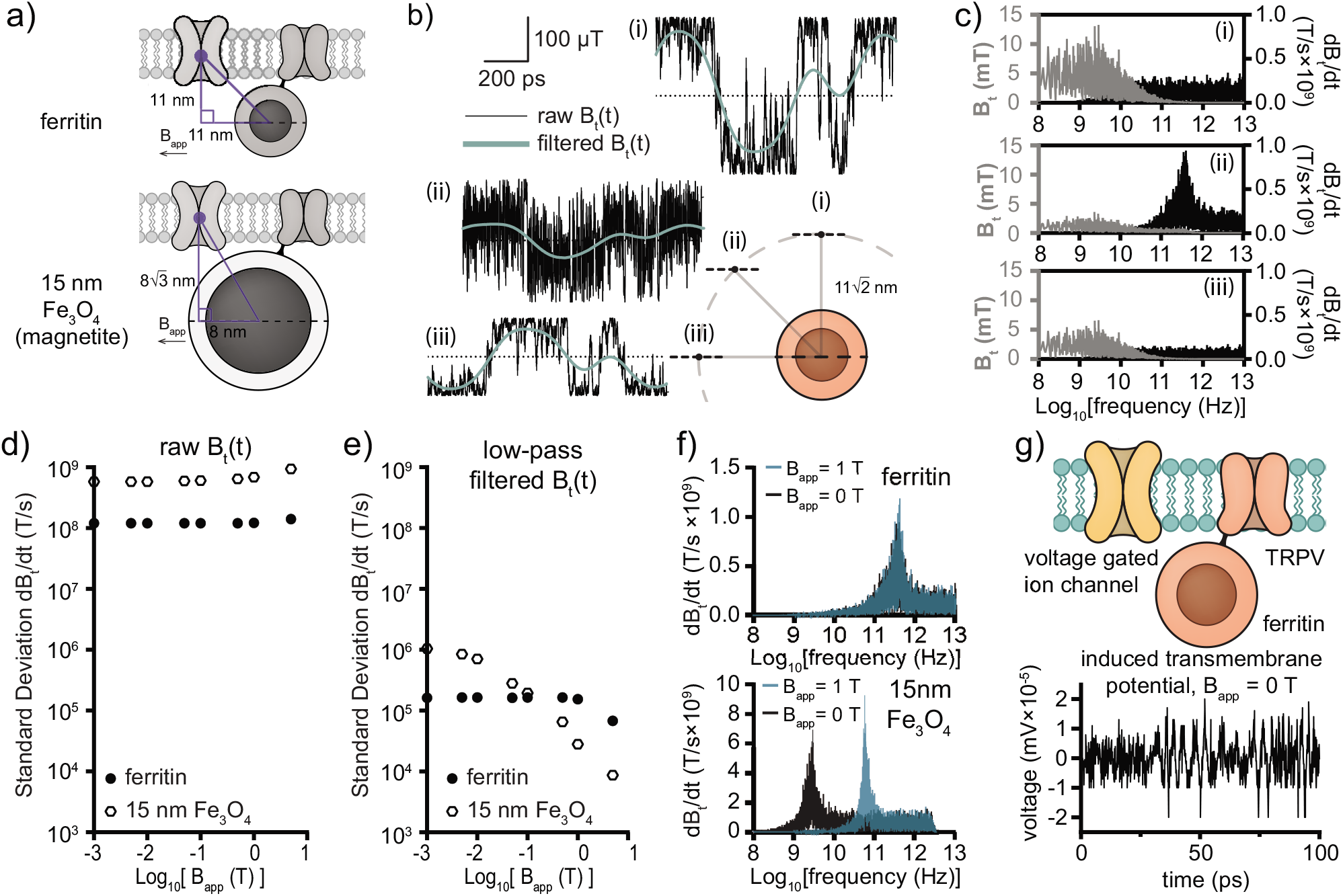
Testing the hypothesis that ferritin could influence neighboring voltage gated ion channels with induced electric fields. a) Simplified geometric assumptions for calculating the component of the field tangential to the membrane are shown explicitly for both a ferritin and a hypothetical 15 nm magnetite particle. b) 1 ns intervals of the tangential projection of magnetic field versus time at three positions equidistant (15.6 nm) from a simulated ferritin. A low pass filter set at 10 GHz reveals signal resulting from stochastic reversal. The dashed line atop the ferritin indicates the orientation of its easy axis. c) Frequency domain representations of *B* (grey) and *dB/dt* (black) at the three locations (i)-(iii) from d) are shown using the full duration of the simulation (6×10^6^ steps, approx. 233 ns) d) Standard deviation of tangential *dB/dt* for simulations of 6×10^6^ steps each (233 ns for ferritin, 397 ns for magnetite) are shown for a range of applied quasimagnetostatic fields. *dB/dt* is dominated by the influence of precession. e) A low pass filter set at 10 GHz for ferritin and 1 GHz for magnetite was applied to the same data represented in panel d), isolating the applied field dependence of dB/dt originating from stochastic reversal. f) A frequency space representation of the *dB/dt* signal from ferritin at no applied field and 1 T shows limited sensitivity to the applied field. A frequency space representation of *dB/dt* from the 15 nm magnetite particle at 0 T and 1 T shows a large influence of the applied field on precession frequency, though not on overall *dB/dt* magnitude. g) The transmembrane voltage predicted from electric fields induced by ferritin is shown for 100 ps of dynamical behavior predicted by the sLLG model. The magnitude of this transmembrane voltage, considered at the center of a neighboring voltage gated ion channel, is too weak to plausibly perturb the gating charge.

To provide a basis of comparison, a similar analysis was performed with a 15 nm magnetite MNP situated on the neuronal membrane in the vicinity of a voltage gated channel (Fig. 3a lower). As a result of its rapid precession, ferritin surprisingly manages to generate a *dB/dt* signal within an order of magnitude compared to this magnetite particle (Fig. 3d). Applying a low pass filter to the signal, the contributions to *dB/dt* from stochastic reversal of the moment are found to be three orders of magnitude weaker (Fig. 3e). These lower frequency components are suppressed in both cases as the applied field increases. Because its exceedingly weak moment and low temperature blocking behavior suggest a strong *H_k_* in the model, a frequency space representation of the *dB/dt* signal from ferritin reveals that its precession frequency does not depend strongly on applied fields below the scale of 1 T (Fig. 3f upper). By contrast, the precession frequency of the magnetite increases more than an order of magnitude under an applied field of 1 T (Fig. 3f lower). The fact that the magnitude of the unfiltered *dB/dt* signal does not increase proportionally with the precession frequency demonstrates a competitive effect between more rapid precession and a magnetic moment more constrained to align with the applied field. Both factors are important in ultimately determining the *dB/dt* experienced by the voltage gated ion channel.

Finally, using this model of the dynamic behavior of the moment of ferritin, we considered the voltage induced across the neuronal membrane at the location of the voltage gated ion channel. The voltage signal is found to be on the scale of 10^−5^ mV, far too small to supply a significant perturbation to the gating charge of the voltage gated ion channel, which would require 10s of mV (Fig. 3f upper). The additive *dB/dt* signal arising from multiple equidistant noninteracting ferritins increases with the square root of their number (Supplementary Section S3). This effectively rules out the possibility that the combined effect of neighboring ferritins could increase this value by many orders of magnitude. Moreover, even if the magntiude of the perturbative membrane voltage had been far larger, it is not clear whether its frequency in the 100s of GHz would have allowed it to influence the conformation of the voltage gated ion channel. Despite the appealing logical features of ferritin acting as source of inductive perturbation on the neuronal membrane for magnetogenetics, our analysis suggests that this hypothesis can be definitively ruled out.

### C. Induced electric fields on ferritin’s mineral core

While ferritin may not act appreciably on the gating charges of nearby channel proteins, local induced electric fields may still be relevant to its indirect mechanistic role in magnetogenetic stimulation. Ferritin is primarily an iron storage protein and recent evidence strongly suggests that the release of Fe^2+^ ions from its core and subsequent oxidation of lipids are likely involved in the indirect stimulation of TRPV channel proteins (Fig. 4a) [33, 34, 45]. This suggested mechanism would be more satisfactory if the role of magnetic fields in this process could be elucidated. Using our model of ferritin, we additionally considered induced electric fields in the direct vicinity of its mineral core, since this is where iron ion release should occur. One additional aspect to consider for the induced fields so close to the center of ferritin is the distribution of magnetization within its mineral core. The magnetic properties of ferritin have variously been attributed to uncompensated surface spins or to the existence of internal magnetic phases [37, 39]. To account for these possibilities, we considered the case of magnetization confined to the outermost layer (“magnetized shell”) or to a small central region (“magnetized center”) in addition to the assumption of uniform magnetization (Fig. 4b). Because precession had been found to dominate the inductive behavior of ferritin (Fig. 3d-f), the electric field was simulated for stable precession at the dominant frequency of 400 GHz predicted in Fig. 3f.

**FIG. 4.**
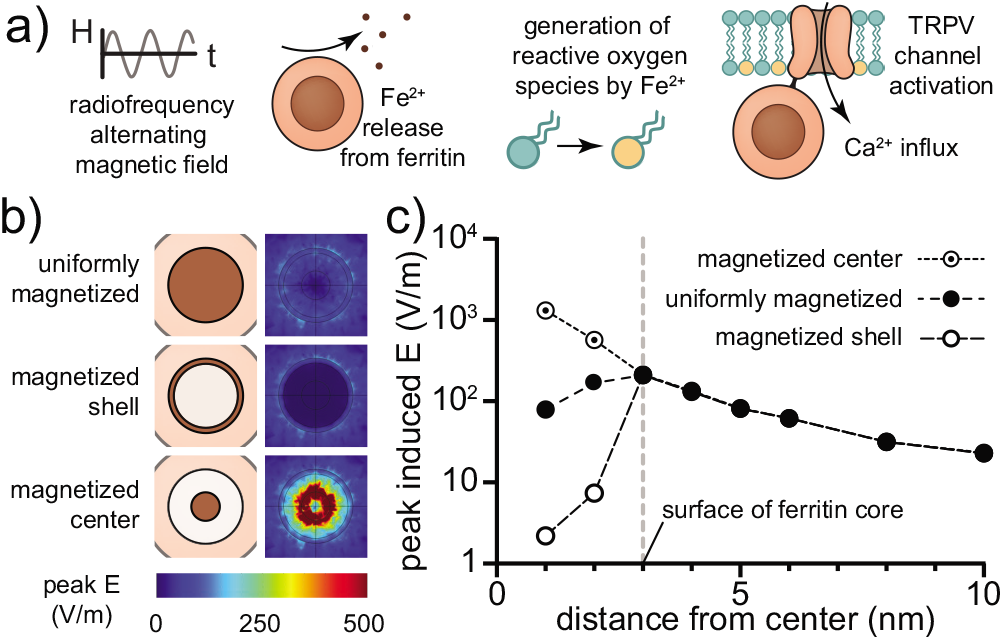
Considering the possible role of electric fields induced at the surface of ferritin in Fe ion release. a) Recent studies have suggested that alternating magnetic fields trigger the release of Fe^2+^ from ferritin, leading to local lipid oxidation of the membrane that indirectly stimulates the channel protein. b) Induced electric fields are considered for stable precession at 400 GHz (see Fig. 3f) assuming different distributions of magnetization within ferritin, including uniform distribution, uncompensated surface spins, or a more strongly magnetized inclusion. c) Variation of the magnitude of the peak induced electric field magnitude as a function of distance from the center of ferritin is plotted for the same three cases. At the surface of ferritin and outside, all three converge to similar values.

The magnitude of the expected induced electric fields converges at the surface of the particle for each of the three cases of magnetization distribution that were considered and similarly diminishes with distance (Fig. 4c). On the surface, the maximum magnitude of this induced electric field is predicted to be about 210 V/m. The induced electric fields inside the particles differ markedly for the different magnetization distributions, with the induced field predicted for the magnetized center exceeding 1000 V/m, and the field for the magnetized shell being suppressed to 10s of V/m. If indeed a small region of increased magnetization occurred in ferritin, it would likely not be confined to its geometric center and thus the electric field realized on the surface could be modestly elevated.

Because ferritin is predicted to generate similar induced electric fields regardless of whether a magnetic field of up to 1 T is applied (Fig. 3d, f), this effect should not be expected to directly liberate Fe ions from the core. Indeed, the predicted electric field magnitude is far less than the electric fields produced by magnetoelectric composite nanoparticles capable of directly triggering chemical reactions, which have been estimated to be as high as 10^7^ V/m [46]. In the case of magnetogenetics, the roles of both ferritin and the externally applied magnetic field are likely to be more subtle than directly breaking chemical bonds, perhaps instead related to mass transport of Fe ions. For instance, the Lorenz force is known to produce observable effects in chemical processes such as electrodeposition or bioelectrocatalytic reactions that are exposed to constant magnetic fields [47, 48]. If ferritin acts as a source of chemically labile iron that is limited by mass transport, externally applied magnetic fields may drive ionic eddy currents that increase Fe ion release. In such a mechanism, the weak electric fields induced by ferritin could perhaps locally enhance the diffusivity of ions at the surface of the mineral core and thus play some role in the overall kinetics of Fe ion release.

## IV. NANOSCALE INDUCTION IN ELECTROCHEMICAL CATALYSIS

### A. Unclear mechanism for catalysis enhanced by alternating magnetic fields

Nanoparticles of various compositions and structures have been vigorously studied as heterogenous catalysts to improve the efficiency of a wide range of chemical processes. This includes instances of metallic single domain ferromagnetic nanoparticles, which can be coupled to externally applied AMFs, for instance to use heat dissipation to control the bulk temperature of a reactor [49]. In a particularly intriguing recent example, an AMF was used to modulate the catalytic enhancement of hydrolysis by iron carbide single domain MNPs coated with a thin layer of nickel that were introduced onto carbon fiber electrodes (Fig. 5a). When an AMF was applied, the presence of these catalytic MNPs was found to reduce the required electrochemical overpotentials by 250 mV at the oxygen evolving electrode and 150 mV at the hydrogen evolving electrode, an enhancement comparable to operating the reaction at approximately 200 °C [10]. Since this effect could not be explained by bulk temperature changes of the reaction solution or other factors, the authors of the study attributed the effect to nanoscale heating of the catalytic MNPs. However, in a subsequent study of chemically similar MNPs monitored by x-ray diffraction while exposed to an AMF, the temperature inside the MNPs matched the bulk temperature within the limits of instrumental uncertainty [13]. Although one interpretation could be that the relevant effects arise from spikes of instantaneous heat flux rather than the establishment of a continuous nanoscale temperature gradient, this serves as yet another striking example in which MNPs have shown clear promise as nanoscale actuators despite an unclear mechanism of action.

**FIG. 5.**
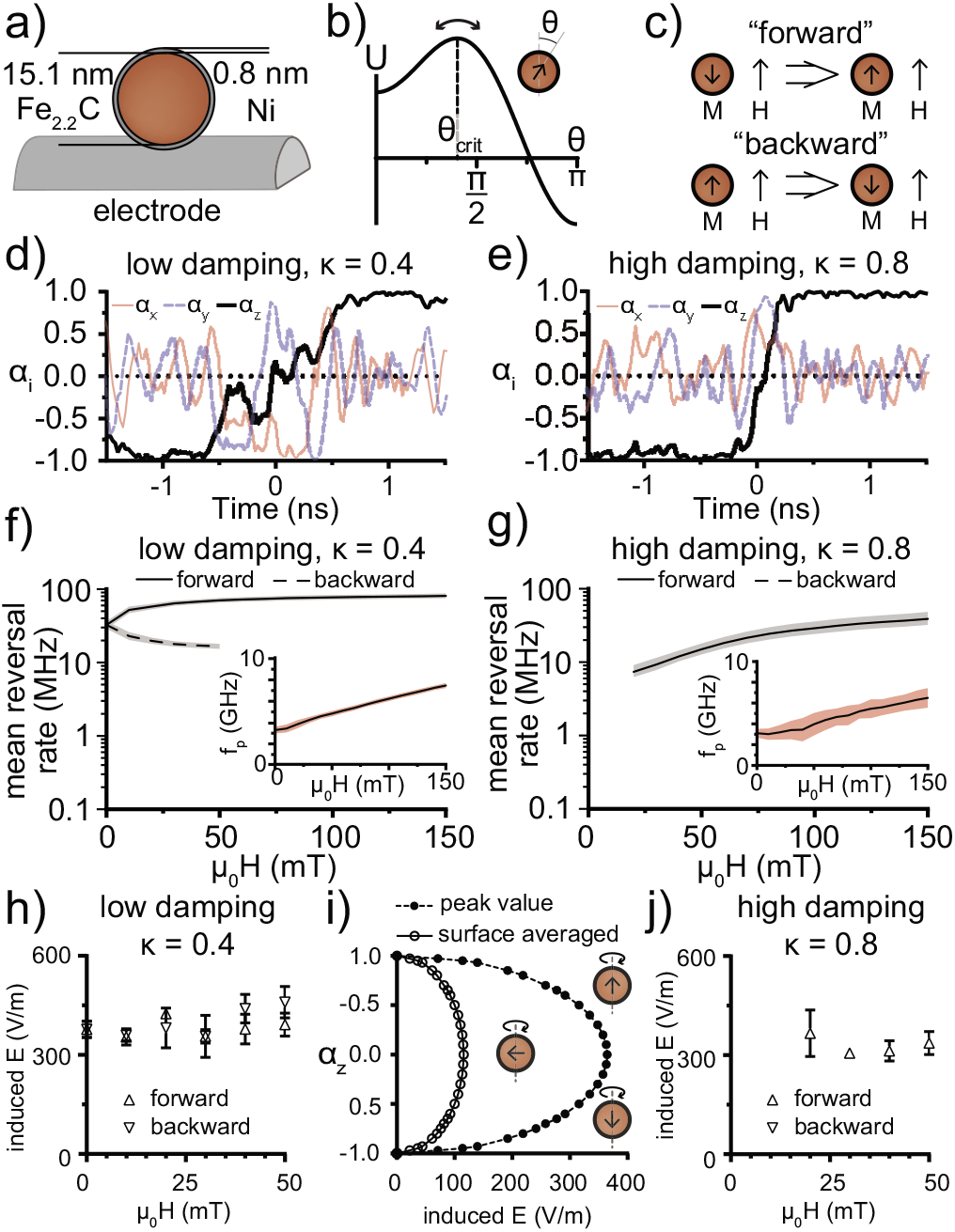
Examining induced electric fields on the surface of catalytic magnetic nanoparticles undergoing hysteresis. a) Assumed dimensions and composition of the magnetic nanoparticles under study. b) The initial orientation of the magnetic moment was set to match the critical angle *θ_crit_*. at which combined energy *U* is maximized, so that the moments proceeded at random to either of the two minima. c) Observed reversal events in which the moment jumped over the barrier were categorized as “forward” if they brought the moment into parallel alignment with applied field and “backward” if they brought the moment into antiparallel alignment. A threshold of *α_z_* = ±0.9 was set to identify reversal events. Representative examples of reversal events are shown for a d) “low damping” case for which *κ* = 0.4 and e) a high damping case for which *κ* = 0.8. The inverse of the time elapsed since the previous reversal is interpreted as an effective reversal rate. f) The mean of all reversal rates observed during 40 separate simulations of 10^4^ steps (230 ns) is shown for the low damping case, sorting reversals according to whether they are forward or backward. The shaded region represents a 95% confidence interval for the mean as determined by empirical bootstrapping with 10^4^ repetitions. The inset shows mean precession frequency *f_p_* extracted from the maxima of FFTs of off axis components of the magnetization for each of the full simulations. Shading corresponds to standard deviation (*N* = 40 for each point). g) The same is shown for the high damping case. Backward reversals were not observed. h) The peak electric field induced at the surface of the catalytic magnetic nanoparticle during representative reversal events was calculated for randomly selected reversal events in the low damping case. Error bars correspond to standard deviation, *N* = 3. i) The peak electric field at the surface of the nanoparticle undergoing stable precession at 4.4 GHz is shown as a function of the orientation of the moment, expressed as its projection *α_z_* along the axis of precession. j) Analogously to h), the peak electric field is plotted for randomly selected reversals with high damping.

Here, we use our computational model of the dynamical motion of the moments of single domain MNPs to consider whether nanoscale electric fields produced by catalytic MNPs could be modulated by the presence of an AMF. One motivation for studying this case is that the reaction in question is electrochemical, providing a more plausible role for nanoscale induced electric fields occurring at the charged interface between the electrode and the solution. Further, it is useful to extend discussion of nanoscale induction to an MNP that is subject to less rapid thermal fluctuation than ferritin and that interacts more robustly with applied fields, since this more closely resembles the majority of cases in which nanoscale effects have been attributed to nanoscale heating. Another reason for selecting the MNPs reported in that particular study of hydrolysis enhancement is that the authors have made sufficient physical characterization data available to select realistic input parameters for modelling [50].

### B. Alternating magnetic fields modulate nanoscale inductive effects in catalytic nanoparticles

Making use of the sLLG model described in Section II, spontaneous reversal events were simulated at constant applied field magnitudes. Although AMFs in the relevant work varied over timescales of μs, this is considerably slower than the relevant features of the dynamical behavior of the single domain MNP moments and it is convenient to consider the influence of the field in its quasistatic limit. Simulations of 10^4^ steps spanned a duration of approximately 200 ns, 2 percent of a full cycle of a field alternating at 100 kHz. The magnitude of the moment of the particle was estimated to be 3.6× 10^5^ *μ_B_* and the *H_k_* was estimated to be 124kA/m, values extracted from previously reported supplementary characterization data [50].

Unlike MNPs consisting of magnetite or other electrically insulating magnetic materials, the effective damping in a metallic nanoparticle is expected to be larger due to eddy currents within its interior [51]. For the purpose of modelling, two sets of assumptions were separately considered—a “low damping” case of *κ* = 0.4 and a “high damping” case of *κ* = 0.8. By applying an initial orientation of the moment at the critical angle corresponding to the top of the energy barrier separating the easy axes, magnetic moments could stochastically progress toward either of the energy minima at *θ* = π or *θ* = 0 (Fig. 5b). Beginning each simulation at the critical angle was intended to sample behavior from each of the minima equally, and the initial transition was excluded from further analysis. The remaining time steps were monitored for further spontaneous transitions over the barrier between the minima. A transition bringing a moment into alignment with the applied field was classified as a “forward” transition (Fig. 5c). Conversely, a transition bringing a moment out of alignment with the applied field was considered a “backward” transition. Representative examples of the time evolution of magnetic moments during selected reversals are shown for low damping (Fig. 5d) and high damping (Fig. 5e). As before, the motion of the moment consists of precession and stochastic jumps over the barrier, albeit much less frequent than for the model of ferritin.

The inverse of the time elapsed since a previous transition can be interpreted as a mean rate of reversal, and this quantity was calculated for both damping conditions over a range of applied field amplitudes. A minimum of 40 simulations of 10^4^ steps were performed at each applied field value from −150 mT to 150 mT in steps of 10 mT, with some simulations capturing multiple reversals. The choice of the damping parameter strongly influences the behavior of the magnetic moment. Low damping leads to comparatively rapid stochastic reversal (frequent forward and backward transitions) that is only suppressed by the application of a sufficient field (Fig. 5f). By contrast, higher damping suppresses stochastic fluctuation over the barrier in the absence of an applied field and limits spontaneous reversal over the barrier at low applied fields (Fig. 5g). Note that points corresponding to backward reversal and forward reversal at low applied fields are not plotted because reversal under these conditions was not observable in the higher damping case at the timescale of the stimulations. The frequency of precession, extracted from fast Fourier transforms of the off-axis components of ***α***, increases linearly with applied field (Fig. 5f and 5g inset). Precession is again expected to dominate the behavior of the nanoscale induced electric fields, and is relatively constant over the 0 to 50 mT range relevant to the application case. The increased standard deviation of the extracted precession frequency for the higher damping case is best explained by greater confinement of the moment to the energy minima, resulting in smaller and noisier off-axis signals. The overall reversal behavior predicted for the higher damping case is more consistent with the available experimental data for hysteretic heat dissipation as a function of applied field amplitude, which suggests effective trapping of magnetic moments below about 15 mT [50].

Finally, electric fields in the vicinity of these catalytic MNPs were modelled based on a small representative sample of the predicted dynamical behavior during reversal events. For the low damping case, the maximum value of the induced electric field predicted at the surface is about 300 to 400 V/m, regardless of the direction of reversal (Fig. 5h). A slight upward trend in this value is observed, which is consistent with the gradual upward trend in precession frequency predicted in the inset of Fig. 5f. It is crucial to recognize that nanoscale inductive effects can be anticipated to depend substantially not only on the frequency, moment, and size of a MNP, but also on the instantaneous angle between the moment and the axis of precession. Fig. 5i shows the calculated induced electric field at the surface of MNPs for cases of pure precessional motion at fixed *α_z_* values, including both the peak electric field magnitude and the surface averaged electric field magnitude. As might be anticipated from symmetry, the induced electric field vanishes when *α_z_* = 1, −1 and is maximized when *α_z_* =0. The peak electric field values obtained during sampled forward reversal events in the high damping case can be shown to be comparable to the low damping case, differing by a small factor that reflects the influence of damping on precession frequency (Fig. 5j).

Taken together, these results suggest that for MNPs capable of effectively thermally trapping moments, the magnitude and extent of induced electric fields are maximized during reversal events. By this reasoning, the generation of these induced electric fields in response to an AMF would be correlated with hysteretic heat dissipation to a degree that would make these effects difficult to distinguish, since effective trapping of moments during the cycle of the AMF is a prerequisite for efficient hysteresis heating via coherent reversal [52]. The peak electric field magnitude was on the order of several hundred V/m, many orders of magnitude weaker than the electric fields existing inside the charged interface that forms on the electrodes and would not be sufficient to trigger electrochemical reactions in isolation. Nevertheless, these fields are far stronger than the initial electric field produced by the electrode that drives the comparatively slow accumulation of charge at its surface when a potential is first applied [53]. Moreover, the direction of the peak electric field induced at the surface of the catalytic MNP is tangential to its surface, with the possibility of introducing electric field components orthogonal to the ones associated with interfacial electrochemical charge accumulation. Thus, either by driving transport within the interface or perturbing the electric field, interaction of these nanoscale induced electric fields with the layers of charges formed at the interface could plausibly influence the process of hydrolysis. The extent of this influence would require modelling of a different sort than the one developed here, and we emphasize that our conclusions are confined to the idea that induced electric fields in the vicinity of MNPs can be correlated with reversal events triggered by AMFs for MNPs with suitable properties for hysteresis heating.

## V. CONCLUSION

This work analyzed the external induced electric fields originating from a frequently overlooked aspect of the expected behavior of magnetic nanoparticles emergent at the nanoscale—precession subject to fluctuation of coherent single domains. In particular, we focused on two case studies in which MNPs have shown promise as nanoscale actuators despite unclear or controversial mechanisms of action. In the first case examined, a role was considered for locally induced electric fields generated by the fluctuation and precession of the magnetic moment of ferritin. Despite the surprising result that ferritin may produce electric fields comparable to a larger magnetite particle, appreciable inductive coupling to voltage gated channel proteins as a source of perturbation was ruled out as a plausible explanation for magnetogenetic stimulation. Nevertheless, the electric fields induced at the surface of ferritin may still be relevant to the mass transport and release of Fe^2+^ that has been suggested to be mechanistically involved. Applying the same model to a separate case involving catalysis of an electrochemical reaction, we considered the possibility that induced electric fields rather than anomalous nanoscale heating might help to explain the enhancement of hydrolysis by electrodes incorporating MNPs in the presence of an AMF. We found that, for MNPs suited to thermally trap magnetic moments, the induction of nanoscale electric fields is expected to be correlated with reversal events triggered by AMFs.

Because magnetic precession was the dominant influence in determining the character of the local induced field, ferromagnetic resonance may offer the most direct means to experimentally test these ideas. For instance, if nanoscale induced electric fields are indeed responsible for enhancing hydrolysis, it may be worthwhile to repeat that experiment under conditions of ferromagnetic resonance to determine whether the enhancement of hydrolysis could be matched or exceeded. Driving resonant precession of the MNPs could have the further technological advantage of extending the duty cycle of the effect, which is only expected to be intermittent in the presence of an AMF. Similarly, the validity of the model of ferritin could potentially be probed with measurements of ferromagnetic resonance, and the relevance of magnetic precession on Fe ion release or local ionic diffusion could be studied empirically, though the weak magnetic properties of ferritin and very high predicted resonance frequency suggest that such studies could be challenging.

The scale of the fields arising from nanoscale induction was consistently estimated to be comparatively weak, at most on the order of 10^3^ V/m for the specific examples considered here. For perspective, Fig. 6 compares the scale of the electric fields involved in a range of technologically relevant phenomena. These values are shown in conjunction with estimates of the relevant timescales at which these fields typically develop and vary, with detailed justification provided in Supplementary Section S4. The characteristic frequencies of nanoscale induction effects tend to be higher than most of the other examples, a feature that is unsurprising since this induction is dominated by their magnetic precession in the GHz frequency range. For MNPs acting as actuators, the effects attributed to nanoscale heating are often subtle and frequently associated with physical rather than chemical changes, such as liposomal release or influence over the surrounding protein corona. Both lipids and proteins notably bear charged groups, and in such cases, relatively weak yet rapidly varying electric fields produced in the immediate vicinity of MNPs may offer explanatory value.

**FIG. 6.**
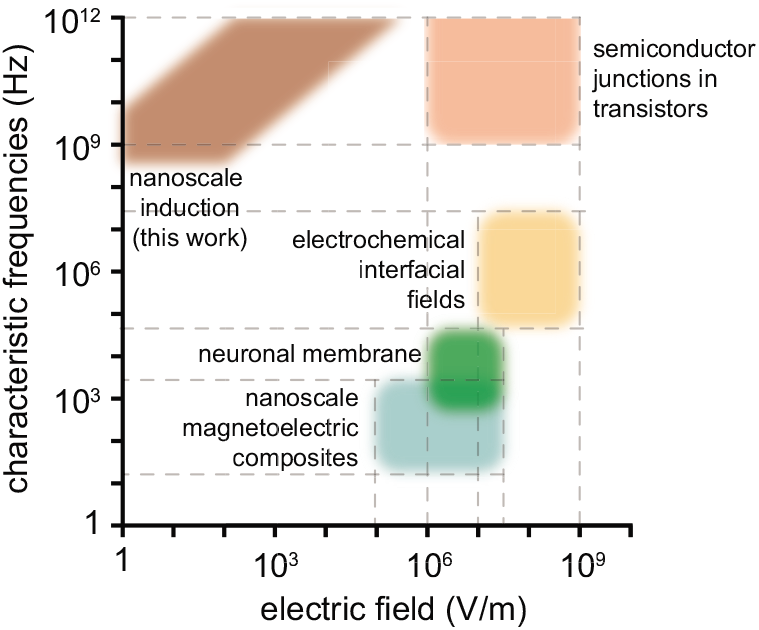
To provide context, the range of expected electric field magnitudes and characteristic frequencies of variation are shown for a variety of technologically relevant areas. The nanoscale induction bounds represented here are based on the generalized parameter space in Fig. 2g. Reasoning behind the other phenomena is explained in Supplementary Section S4.

If nanoscale induced electric fields do indeed play an unappreciated role in localized actuation effects of MNPs, it is worthwhile to consider how single domain MNPs could be engineered to deliberately maximize induced fields at their surfaces. Fig. 2g captures the essential parameter space, showing induced electric field normalized to precession frequency in GHz as a function of particle size and magnetization. Since the expected induced field magnitude increases with size, magnetization, and precession frequency, one strategy could be to design MNPs that maximize these quantities. For instance, selecting a metal or metal alloy that has high magnetocrystalline anisotropy would allow larger particles to support a single domain state and would increase the effective *H_k_*, which is predicted by the sLLG equation to raise the frequency of precession. In a biomedical setting, the limited penetration depth of stimuli in the 10s of GHz or higher would likely preclude these MNPs from being driven at resonance. A somewhat counterintuitive alternative strategy could instead be to use relatively large oxide nanoparticles (on the order of 100 nm diameter) with low anisotropy and moderate to low saturation magnetization. This would permit ferromagnetic resonance to be driven at lower frequencies, which might offer the benefit of greater physiological penetration depth of the magnetic stimulus. The intrinsically lower damping expected for oxides due to the lack of eddy currents might also result in sharper resonance and more efficient energy transfer. One challenge that could be anticipated for enlarged nanoparticles is the eventual development of spin wave modes in which the net moment of the MNP is partially cancelled.

The number and variety of investigations triggering or enhancing biological and chemical processes in a localized fashion with magnetic nanoparticles has grown substantially in recent years and seems poised to continue. While it may be intuitive to conceptualize the influence of MNPs as a scaled down analogue of our macroscopic experience such as nanoscale heating, alternative explanations must be explored when such assumptions are contradicted by both theory and experiment. The local induced electric fields that we have considered here rely on coherent precession and reversal of single domains, making them an emergent phenomenon of nanoscale magnetic materials. Gaining a clearer understanding of the mechanism of nanoscale actuation in these systems, whether through the effect we have examined or through some other influence yet to be identified, is essential for optimizing these systems and envisioning novel techniques for nanoscale actuation.

## ACKNOWLEDGMENTS

M.G.C. was supported by an ETH Zurich Postdoctoral Fellowship. S.S. is supported by the Branco Weiss Fellowship of the Society in Science. Sonia Monti assisted with illustrations.

