## Supplementary for "Localized nanoscale induction by single domain magnetic particles"

### Supplemental Material for “Localized nanoscale induction by single domain magnetic particles”

**Supplementary Section S1: Physical properties assumed in electric field simulations**

The methods used to model the electric field induced by fluctuations of the moments of magnetic nanoparticles are described primarily in Section II of the main text. For clarity, here we list the material properties assumed for the various domains of the finite element model implemented in COMSOL. Reports of these values vary widely and depend on frequency, so our assumed quantities are representative based on a broad sampling of relevant literature [1–8]. For tissue, it was assumed that $\mu_r \approx 1$.

**Table S1.** Assumed electrical properties for finite element modelling at various frequencies

| Freq. (GHz) | Extracellular | | Membrane | | Cytoplasm | |
|---|---|---|---|---|---|---|
| | conductivity (S/m) | relative permittivity | conductivity (S/m) | relative permittivity | conductivity (S/m) | relative permittivity |
| 1 | 5.00E-01 | 1.50E+02 | 3.00E-07 | 1.20E+01 | 3.00E-01 | 1.00E+02 |
| 10 | 2.00E+00 | 1.17E+02 | 1.89E-05 | 8.33E+00 | 3.53E+00 | 7.00E+01 |
| 100 | 3.50E+00 | 8.33E+01 | 3.74E-05 | 4.67E+00 | 6.77E+00 | 4.00E+01 |
| 1000 | 5.00E+00 | 5.00E+01 | 5.60E-05 | 1.00E+00 | 1.00E+01 | 1.00E+01 |

**Supplementary Section S2: Details of Implementation for Numerical Solution and Estimating $H_k$**

Picking an appropriate timestep for solving the stochastic Landau-Lifshitz-Gilbert equation relies on having a sufficient number of steps per period of precession. The maximum expected rate of precession is related to the scale of the fields (internal and applied) that the moment will experience. The frequency of Larmor precession $f_p$ for electrons in a field $H$ is as follows:

$$f_p = \mu_0 H \frac{\gamma}{2\pi}. \tag{S1}$$

With this information, time can be scaled by a factor that leaves the differential equation unitless and ensures that when $t_p = 1$, $t = \frac{1}{f_p}$, i.e. the period of precession. Thus,

$$t_p = t\, \mu_0 H \frac{\gamma}{2\pi}. \tag{S2}$$

For ferritin, over the range of applied fields considered (50 µT to 5 T), $H_{ap} < H_k$, such that the anisotropy field will determine the maximal precession frequency and Equation 1 of the main text can be rescaled as follows:

$$\frac{d\vec{\alpha}}{dt_p} = 2\pi\left[-\frac{1}{1+\kappa^2}\vec{\alpha}\times\frac{\vec{\mathrm{H}}_{eff}}{H_k} - \frac{\kappa}{1+\kappa^2}\vec{\alpha}\times\left(\vec{\alpha}\times\frac{\vec{\mathrm{H}}_{eff}}{H_k}\right)\right]. \tag{S3}$$

For the magnetite and iron carbide particles considered, at the largest field value of 5 T, $H_{ap} \gg H_k$, such that the applied field will determine the maximum precession frequency. $t_p$ can be defined accordingly:

$$\frac{d\vec{\alpha}}{dt_p} = 2\pi\left[-\frac{1}{1+\kappa^2}\vec{\alpha}\times\frac{\vec{\mathrm{H}}_{eff}}{H_{ap}} - \frac{\kappa}{1+\kappa^2}\vec{\alpha}\times\left(\vec{\alpha}\times\frac{\vec{\mathrm{H}}_{eff}}{H_{ap}}\right)\right]. \tag{S4}$$

To solve these differential equations, we implemented a 4$^{th}$ order Runge-Kutta method in Python. In our algorithm, the i$^{th}$ component of stochastic noise is introduced as a normally distributed random variate multiplied by a constant $H_{th0}$, i.e.,

$$\vec{H}_{th,i}(t) = H_{th0}\, N[0,1]. \tag{S5}$$

$H_{th0}$ depends both on physical parameters supplied by the problem, and on the stepsize $\Delta t$. For our simple model, to ensure convergence to physically motivated behavior, we found $H_{th0}$ empirically. Keeping the time scaling constant for each particle over all relevant simulations, we first removed the influence of anisotropy and considered the time averaged projection of their moments along the field direction for a logarithmic sequence of applied fields between 50 µT and 5 T. The noise was selected to ensure convergence to the Langevin function at points corresponding to $M/M_s = 0.25$ for ferritin and $M/M_s = 0.75$ for magnetite. The application of the noise terms determined in this way to the rest of the range of field values was sued to check the ability of the model to converge to the Langevin function, as noted in the main text. Although some reported attempts to model ferritin show fits to a modified Langevin function with an additional superimposed paramagnetic term [9,10], the contribution to the internal field experienced by the particles is likely small and we have focused on the superparamagnetic component since it is the part most relevant to fluctuation and generation of inductive noise.

**Estimating $H_k$**

Kinetically limited fluctuation of magnetic moments of superparamagnetic particles is well studied, and characterization that directly or indirectly probes reversal behavior enables the estimation of $H_k$ the field at which the effective barrier to reversal vanishes. Following Néel, this reversal behavior is described by the following equation [11]:

$$\tau = \tau_0 \, Exp\left(\frac{U_B}{k_B T}\right). \tag{S6}$$

Here, as noted in the main text, $\tau$ is the timescale of stochastic reversal, $\tau_0$ is a characteristic attempt rate, $U_B$ is the energy barrier to reversal, $k_B$ is the Boltzmann constant, and $T$ is temperature. One particularly thorough analysis of reconstituted horse spleen ferritin characterized at multiple timescales and temperatures found $\tau_0 = 9 \times 10^{-12}$ s and $U_B = 4.39 \times 10^{-21}$ J [9]. Rather than relying on a single report, we have instead made assumptions within the reasonable range of reported values. One way to estimate $U_B$ for ferritin is from its reported blocking temperature, which has often been reported as 40 K [12]. Assuming that at this temperature, the rate of reversal would match the timescale of a slow magnetization measurement, about 10 s,

$$k_B T \ln\left(\frac{\tau}{\tau_0}\right) = U_B = (1.381 \times 10^{-23}\ \mathrm{JK^{-1}})\ (40\ \mathrm{K}) \ln\left(\frac{10\ \mathrm{s}}{10^{-9}\ \mathrm{s}}\right) = 1.3 \times 10^{-20}\ \mathrm{J}. \tag{S7}$$

This implies $\tau \approx 2.2 \times 10^{-8}$s at 300 K for ferritin. Cases similar to the 15 nm magnetite particle are frequently modeled in the context of heat dissipation under alternating fields. The height of the anisotropy barrier is typically related to the anisotropy energy $K$ and the magnetized volume $V_m$,

$$U_B = K_u V_m. \tag{S8}$$

The leading order term of the magnetocrystalline anisotropy of magnetite has cubic symmetry and is negative, $K_c \approx -1.4 * 10^4\ \mathrm{J\ m^{-3}}$ [13], although contributions from shape and surface can play a larger role [14]. To enable straightforward implementation of the stochastic Landau-Lifshitz-Gilbert (LLG) equation, we assume a uniaxial form with $K_u \approx |K_c|$, a common assumption which tends to overestimate the effective barrier. For a 15 nm particle, this implies $U_B = 2.5 \times 10^{-20}$ J, giving a characteristic timescale of reversal of $3.9 \times 10^{-7}$s at 300 K.

As will be shown in the next section, the energy barrier is incorporated into the stochastic LLG equation through the anisotropy field $H_k$ for each case. For ferritin, this can be expressed in terms of $U_B$ and its moment $m$:

$$H_k = \frac{2U_B}{\mu_0 m} = \frac{2(1.3 \times 10^{-20}\ \mathrm{J})}{(4\pi \times 10^{7}\ \mathrm{N\ A^{-2}})(300 * 9.274 \times 10^{-24}\mathrm{J\ T^{-1}})} = 7.3 \times 10^{6}\ \mathrm{A\ m^{-1}}. \tag{S9}$$

This implies that a field of 9.1 T would be needed to make the barrier to reversal vanish for ferritin, a situation mostly attributable to its exceedingly weak moment. For the magnetite particle,

$$H_k = \frac{2K_u}{\mu_0 M_s} = \frac{2(1.4 \times 10^{4}\ \mathrm{J\ m^{-3}})}{(4\pi \times 10^{-7}\ \mathrm{N\ A^{-2}})(4.7 \times 10^{5}\ \mathrm{A\ m^{-1}})} = 4.7 \times 10^{4}\ \mathrm{A\ m^{-1}}. \tag{S10}$$

This ensures that a field of only 59 mT is required to make the barrier to reversal vanish for the magnetite particle.

#### Supplementary Section S3: Combined Effects of Neighboring Ferritin

Because schemes that involve multiple ferritin units in close proximity to one another have been reported [15], we investigated the potential for additive effects. At a point on a line of symmetry equidistant from two ferritins, both filtered and unfiltered $dB/dt$ increase by a factor approaching $\sqrt{2}$, an expected result for the addition of two sources of uncorrelated noise with the same amplitude (Fig. S1 left). The rapid dropoff of the field ensures that only nearest neighbor contributions are most relevant (Fig. S1 right). While it is conceivable that placing several inductive noise sources in close proximity can modestly enhance local effects, it should be emphasized that this does not imply the possibility of collective effects over larger scales.

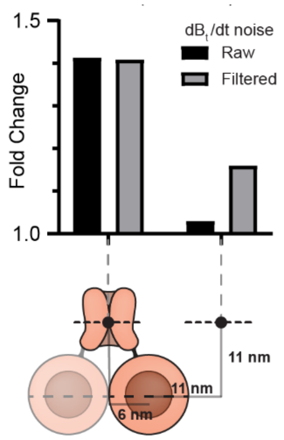


**Fig. S1** Additive effects of magnetic field produced by two adjacent non-interacting ferritins. The fold increase in the standard deviation of the noise signal is shown for two points in the center of the membrane. The first point is equidistant from the two ferritins and the second is 15.6 nm from the closer of the two ferritins.

**Supplementary Section S4: Electric Field Magnitudes and Timescales in Various Phenomena**

| Phenomenon | Characteristic Frequency (Hz) | Justification | Characteristic E (V/m) | Justification |
|---|---|---|---|---|
| Nanoscale inductive effects (this work) | High 100s of MHz to THz | Below the mid-100s of MHz, interfacial capacitive effects might become relevant | Values shown on plot | Plotted from limiting cases of parameter space: 6 nm diameter with Ms = 20 kA/m and 100 nm diameter with Ms = 200 kA/m |
| Action potentials | 100s of Hz to 10s of kHz | Action potentials observed with patch clamping occur over several ms, relevant features varying on the 10s or 100s of μs | $10^6$ to low $10^7$ | The electric field inside the neuronal membrane can be estimated from the potential difference on the order of 10s to 100 mV and approximately 10 nm membrane thickness |
| Transistors | GHz to >1THz | Transistor speed is influenced by many intrinsic and extrinsic factors, however transistors that operate in the low GHz are commercially available and nanoscale transistors can be demonstrated to operate past 1 THz. [16] | $10^6$ to $10^9$ | A wide value is given because the electric field values reached depend on e.g. doping concentrations and band gaps. Electron holography has been used to directly observe these fields. [17] |
| Electrochemical interfacial phenomena | MHz to GHz | In principle, limited by Debye time, with typical values in the ns to μs range. [18] | $10^7$ to $10^9$ | The Gouy-Chapman model of charge accumulation at interfaces in solution predicts electric fields on the order of $10^9$ V/m. Various experimental methods have found similar values. [19] |
| Magnetoelectric composites | 10s of Hz to kHz | In principle, magnetoelectric composites exhibit peak coupling at their mechanical resonance frequency. In practice, frequencies on the order of 10s of Hz to kHz are used even for nanoparticles. | $10^5$ to mid $10^7$ | The upper limit comes from an estimate in a recent report that references similar values found in nanostructures. [20] Neuromodulation was accomplished with similar nanoparticles exhibiting a far lower coupling coefficient, giving the lower bound. [21] |
